# Small molecule analysis of extracellular vesicles produced by *Cryptococcus gattii*: identification of a tripeptide controlling cryptococcal infection in an invertebrate host model

**DOI:** 10.1101/2021.01.18.427068

**Authors:** Flavia C. G. Reis, Jonas H. Costa, Leandro Honorato, Leonardo Nimrichter, Taícia P. Fill, Marcio L. Rodrigues

## Abstract

The small molecule (molecular mass < 900 Daltons) composition of extracellular vesicles (EVs) produced by the pathogenic fungus *Cryptococcus gattii* is unknown, which limits the understanding of the functions of cryptococcal EVs. In this study, we analyzed the composition of small molecules in samples obtained from solid cultures of *C. gattii* by a combination of chromatographic and spectrometric approaches, and untargeted metabolomics. This analysis revealed previously unknown components of EVs, including small peptides with known biological functions in other models. The peptides found in *C. gattii* EVs had their chemical structure validated by chemical approaches and comparison with authentic standards, and their functions tested in a *Galleria mellonella* model of cryptococcal infection. One of the vesicular peptides (isoleucine-proline-isoleucine, Ile-Pro-Ile) improved the survival of *G. mellonella* lethally infected with *C. gattii* or *C. neoformans*. These results indicate that small molecules exported in EVs are biologically active in *Cryptococcus.* Our study is the first to characterize a fungal EV molecule inducing protection, pointing to an immunological potential of extracellular peptides produced by *C. gattii*.

## Introduction

*Cryptococcus gattii* is a fungal pathogen that causes disease in immunocompetent individuals. This fungus was responsible for outbreaks in the Pacific Northwest and in the Vancouver Island (1). *C. gattii* virulent strains, which are endemic in Brazil (2), likely emerged from South America (3). *C. gattii* can cause severe lung disease and death without dissemination. In contrast, its sibling species *C. neoformans* disseminates readily to the central nervous system (CNS) and causes death from meningoencephalitis (1). *C. gattii* and *C. neformans* share major virulence determinants, including the ability to produce extracellular vesicles (EVs) (4–6). EVs are membranous structures produced by prokaryotes and eukaryotes, including fourteen fungal genera (7). In fungi, they were first characterized in culture fluids of *C. neoformans* (6). A decade later, *C. gattii* was also demonstrated to produce EVs in liquid matrices (4).

The perception that EVs are essential players in both physiology and pathogenesis of fungi is now consolidated. Much of the knowledge on the functions of fungal EVs has derived from studies of their composition. During the last decade, proteins, lipids, glycans, and nucleic acids were characterized as components of fungal EVs (8,9). Molecules of low molecular mass, however, have been overlooked. A recent study has characterized the small molecule composition of *Histoplasma capsulatum* (10) and *Penicillium digitatum* EVs (11), but the low molecular mass components of other fungal EVs are unknown. Considering the molecular diversity found in both *H. capsulatum* and *P. digitatum*, it is plausible to predict that many still unknown functions of EV components of low molecular mass remain to be characterized. In fact, the metabolome analysis of *P. digitatum* EVs revealed the presence of phytopathogenic molecules that inhibited the germination of the plant host’s seeds (11).

We have recently described a protocol for the isolation of cryptococcal EVs through which the vesicles were obtained from solid fungal cultures (5). Although the general properties of fungal EVs obtained from solid cultures resembled those described for vesicles obtained from liquid media, a recent analysis of the protein composition of cryptococcal EVs obtained from solid medium revealed important differences in comparison to those obtained in early studies using liquid cultures (12,13). This observation and the fact that culture conditions impact the composition of small molecules in *H. capsulatum* EVs (10) reinforce the importance of the compositional characterization of vesicles obtained from solid medium.

In this manuscript, we characterized the low mass components of EVs produced by *C. gattii*. The synthesis of some of the small molecules detected in the EVs revealed a vesicular peptide that protected an invertebrate host against a lethal challenge with *C. gattii* in a dose-dependent fashion. These results indicate the existence of new venues of exploration of the functions of EVs in fungal pathogens, and suggest that small molecules of fungal EVs have immunological potential.

## Results

### Small molecule characterization of *C. gattii* EVs

*C. gatti* EV samples were prepared as independent triplicates. EV extracts were analyzed by ultra-high performance liquid chromatography-tandem mass spectrometry (UHPLC-MS/MS), and the data submitted to molecular networking analysis in the Global Natural Product Social Molecular Networking (GNPS) platform, an interactive online small molecule–focused tandem mass spectrometry data curation and analysis infrastructure (14). Molecular networking using high-resolution MS/MS spectra allows the organization of vesicular compounds in a visual representation (15,16). In this analysis, each node is labeled by a precursor mass and represents a MS/MS spectrum of a compound, and compounds of the same molecular family are grouped together, connected by arrows, forming clusters of similarity (15–18). Since the molecules can be identified in a database through their fragmentation patterns and are represented in the molecular networking, the benefits of this approach include fast dereplication, identification of similar compounds, and effortless comparisons between different metabolic profiles or conditions (16,17).

The cluster-based molecular networking analysis revealed secondary metabolites present in the *C. gattii* EVs. The molecules detected in our analysis were classified as EV components if they were detected in the three replicates. Using this criterion, our small molecule analysis identified 13 genuine components of the *C. gattii* EV samples (Table 1). This analysis revealed previously unknown components of EVs, including peptides, amino-acids, vitamins, and a carboxylic ester. The metabolites were identified through hits in the GNPS database (Supplemental Figures 1-13) and corresponded to Ile-Pro-Ile (*m/z* 342.2384), Phe-Pro (*m/z* 263.1387), Pyro Glu-Ile (*m/z* 243.1335), Pyro Glu-Pro (*m/z* 227.1022), Leu-Pro (*m/z* 229.1544), Pyro Glu-Phe (*m/z* 277.1180), Val-Leu-Pro-Val-Pro (*m/z* 652.4025), cyclo (Trp-Pro) (*m/z* 284.1393), cyclo (Tyr-Pro) (*m/z* 261.1234), tryptophan (*m/z* 205.0972), asperphenamate (*m/z* 507.2278), riboflavin (*m/z* 377.1456) and pantothenic acid (*m/z* 220.1181). The structures and MS data of the detected metabolites are shown in Figure 1 and Table 1, respectively. The cluster-based molecular networking analysis of the *C. gattii* EV components is detailed in Figure 2.

**Figure 1.**
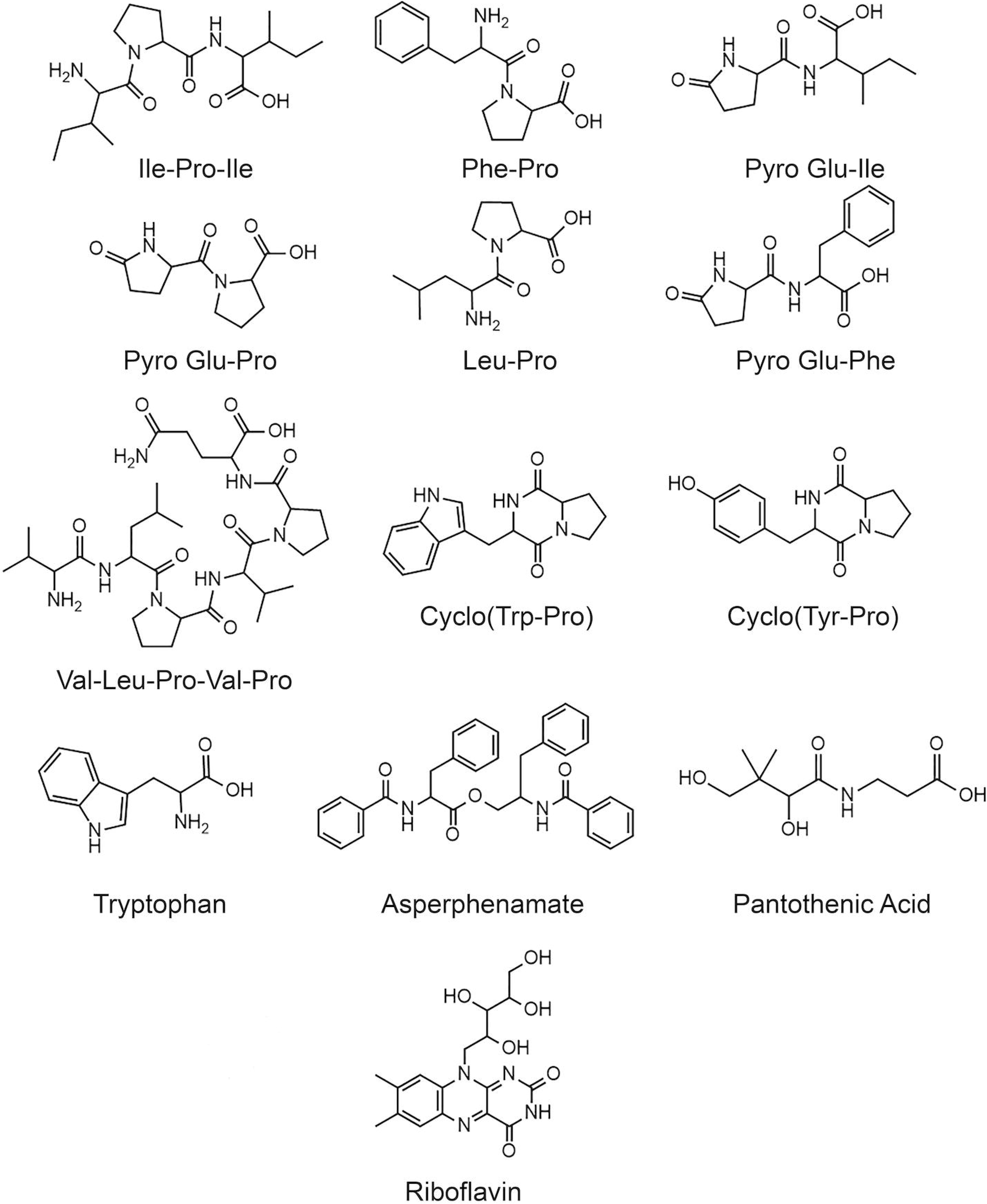
Structures of the metabolites identified in *C. gatti* EVs through the GNPS MS/MS database.

**Table 1.**
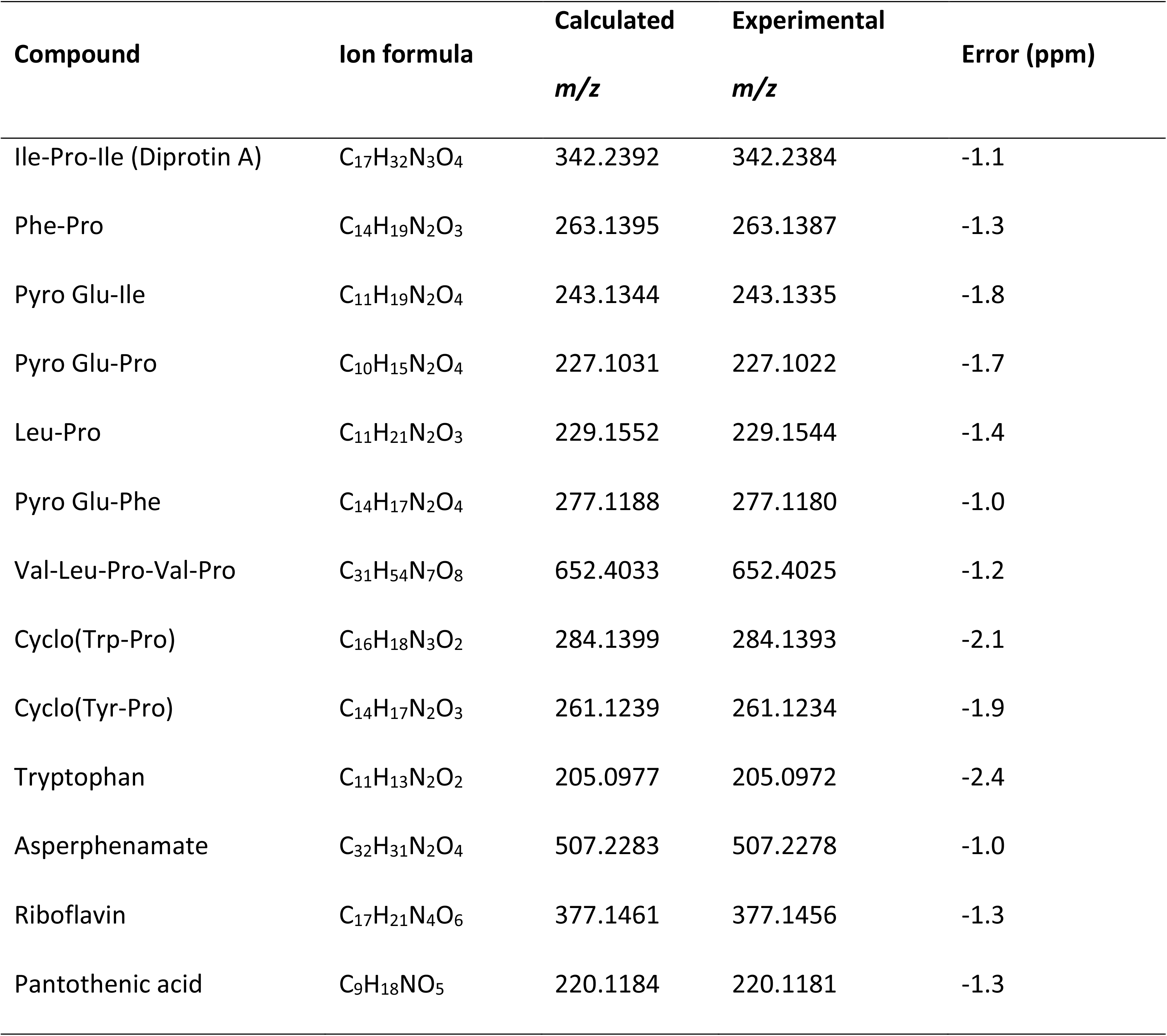
MS data obtained for *Cryptococcus gattii* secondary metabolites detected on EVs.

**Figure 2.**
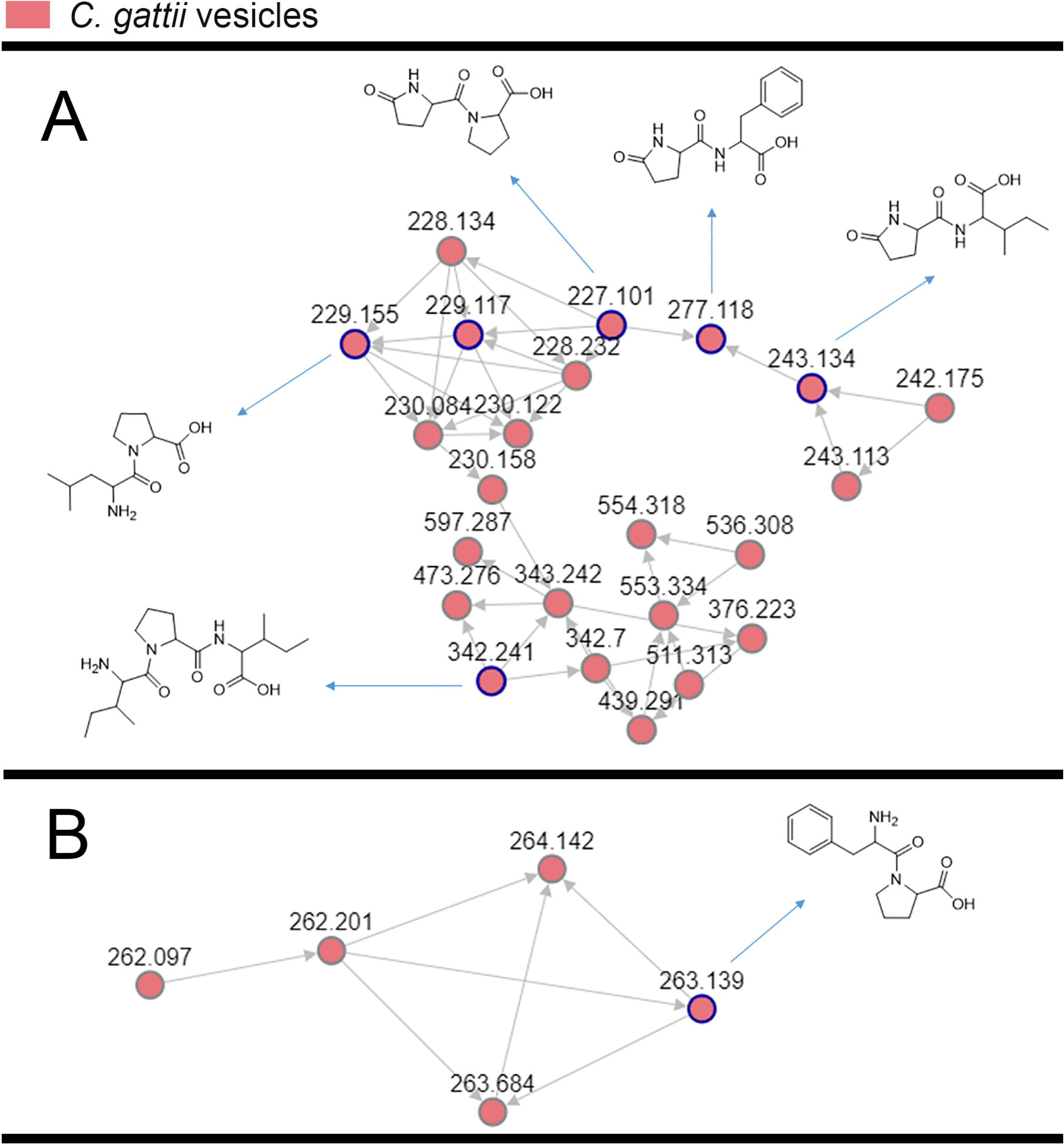
Clusters A and B of the molecular networking obtained for the *C. gattii* EV cargo. Nodes circled in blue indicate molecules identified by comparison with the GNPS platform database. Authentic standards were used to validate the hits within the GNPS database.

For validation of some key GNPS hits, we performed another round of spectrometric characterization of *C. gattii* small molecules including additional criteria as follows. We classified as authentic EV compounds those whose structure was observed in EV extracts, but not in mock samples (extracted from sterile culture medium). Finally, these compounds obligatorily had chromatographic and spectrometric properties similar to those of synthetic standards. Due to the easiness in chemical synthesis and lack of functional information in the literature, the linear dipeptides Phe-Pro, pyro-Glu-Ile, pyro-Glu-Pro, Leu-Pro, and pyro-Glu-Phe, and the tripeptide Ile-Pro-Ile were selected for the validation assays. We then searched for their presence in EV and mock extracts. Six peptides were classified as authentic EV components according to these criteria (Table 2). Indeed, this analysis revealed similar fragmentation patterns and retention times for the vesicle peptides and the standard metabolites (Figure 3). The peptides exhibited typical fragments of protonated amino acids at *m/z* 70.06, 86.09, 116.07 and 120.08 (Figure 4). In compounds containing proline, fragments at *m/z* 116.07 and 70.06 corresponded, respectively, to the loss of protonated proline and subsequent loss of H_2_O and CO. In peptides composed by isoleucine or leucine, fragments at *m/z* 132.02 and 86.09 corresponded, respectively, to protonated leucine/isoleucine and subsequent loss of H_2_O and CO. Finally, the loss of H_2_O and CO in protonated phenylalanine formed the major fragment ion at *m/z* 120.08 (19). Assuming that the vesicular components are synthesized within the cells and exported extracellularly, we also analyzed cellular and supernatant extracts. The six peptides listed in Table 3 were also found in these extracts (data not shown).

**Table 2.** Chromatographic identification of peptides in cryptococcal EVs*. Sample (retention time, min)

| Peptide | Control | Mock | EVs | Synthetic standards |
| --- | --- | --- | --- | --- |
| Ile-Pro-Ile | NF | NF | 4.03 | 3.76 |
| Phe-Pro | NF | NF | 3.18 and 3.64 | 3.28 and 3.62 |
| Pyro-Ile | NF | NF | 3.55 | 3.53 |
| Pyro-Pro | NF | NF | 1.8 | 1.8 |
| Leu-Pro | NF | NF | 2.83 | 2.78 |
| Pyro-Phe | NF | NF | 4 | 4 |

\*Peptide identification was performed in blank samples (control) in addition to preparations obtained from sterile medium (mock) or fungal EVs. The results were compared to those obtained with synthetic peptides. NF, not found.

**Figure 3.**
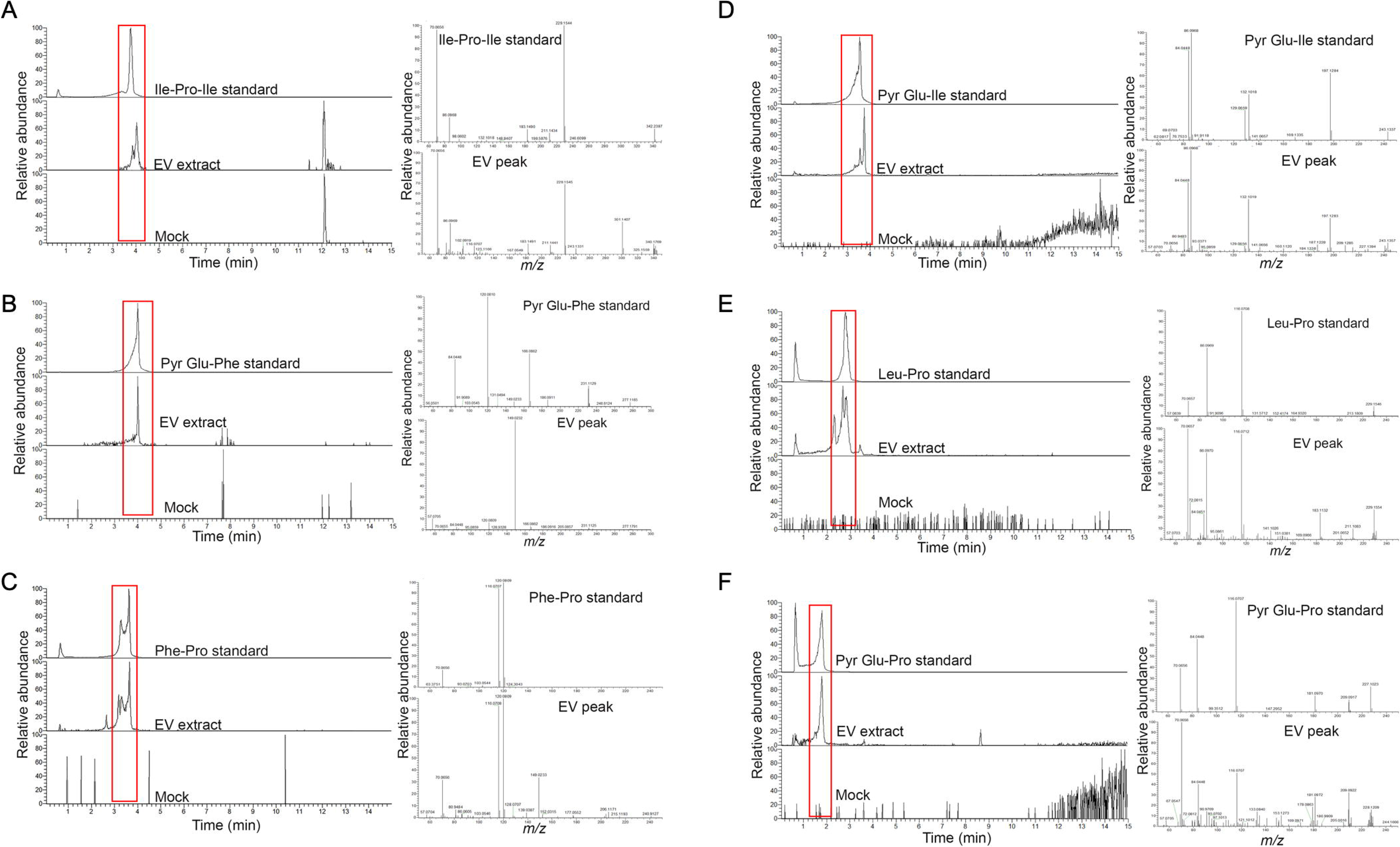
Structural analysis of EV peptides produced by *C. gattii*, including Ile-Pro-Ile (A), pyr-Glu-Phe (B), Phe-Pro (C), pyr-Glu-Ile (D), Leu-Pro (E), and pyr-Glu-Pro (F). For each peptide, the chromatographic separation of synthetic standards, EV extracts, and control (mock) samples is presented on the left side of each panel. The peaks with retention times similar to the corresponding standards (red boxed area) were selected for MS analyses, which are shown on the right side of each panel. These analyses confirmed that the structural match between the EV components and the synthetic standards.

**Figure 4.**
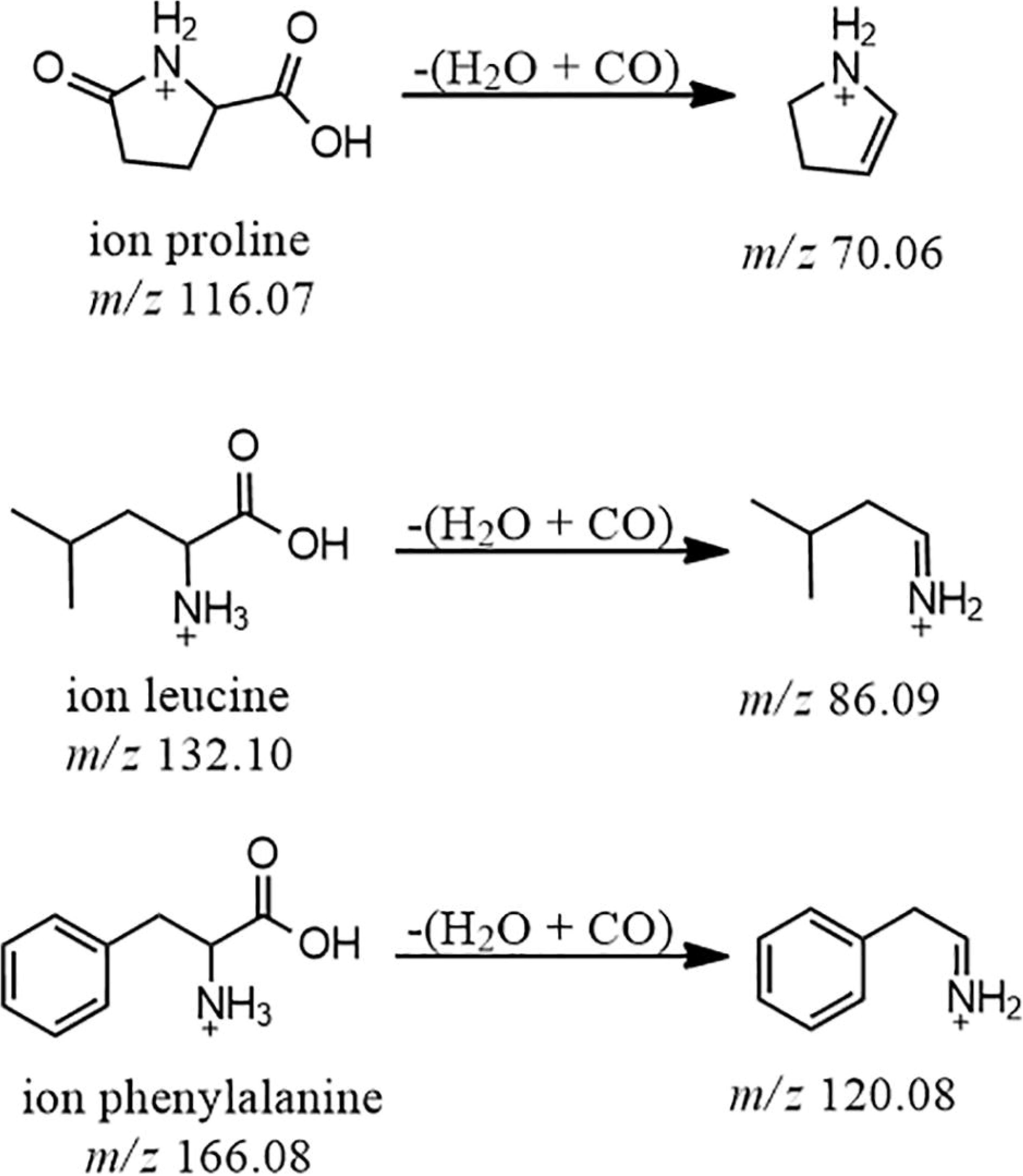
Main mass fragments obtained for amino acids leucine, proline and phenylalanine.

### Biological activity of EV peptides of *C. gattii*

After characterization of Ile-Pro-Ile, Phe-Pro, Pyro-Ile, Pyro-Pro, Leu-Pro, and Pyro-Phe as authentic EV components of *C. gattii*, we used their synthetic forms to analyze their possible biological activities. On the basis of the previously reported ability of fungal peptides to kill bacteria (20), we initially tested their antibacterial capacity against *Staphylococcus aureus* and *Pseudomonas aeruginosa.* None of the peptides had any effect on microbial growth (data not shown). Since cryptococcal EVs regulate intercellular communication (4), we also speculated that the peptides could mediate quorum sensing, Titan cell formation or capsule growth. Once again, none of the peptides had any apparent effects on these processes in *C. gattii* (data not shown).

It has been recently reported that fungal EVs, including cryptococcal vesicles, protect mice and the invertebrate host *Galleria mellonella* against lethal challenges with pathogenic fungi (12,21–23). The vesicular molecules responsible for the protection remained unknown. We then asked whether the peptides listed in Table 3 could protect *G. mellonella* against a lethal challenge with *C. gattii*. We compared the mortality curves of *G. mellonella* infected with *C. gattii* alone with the mortality of the invertebrate host receiving *C. gattii* and each of the peptides at 10 μg/ml (Figure 5A).

**Figure 5.**
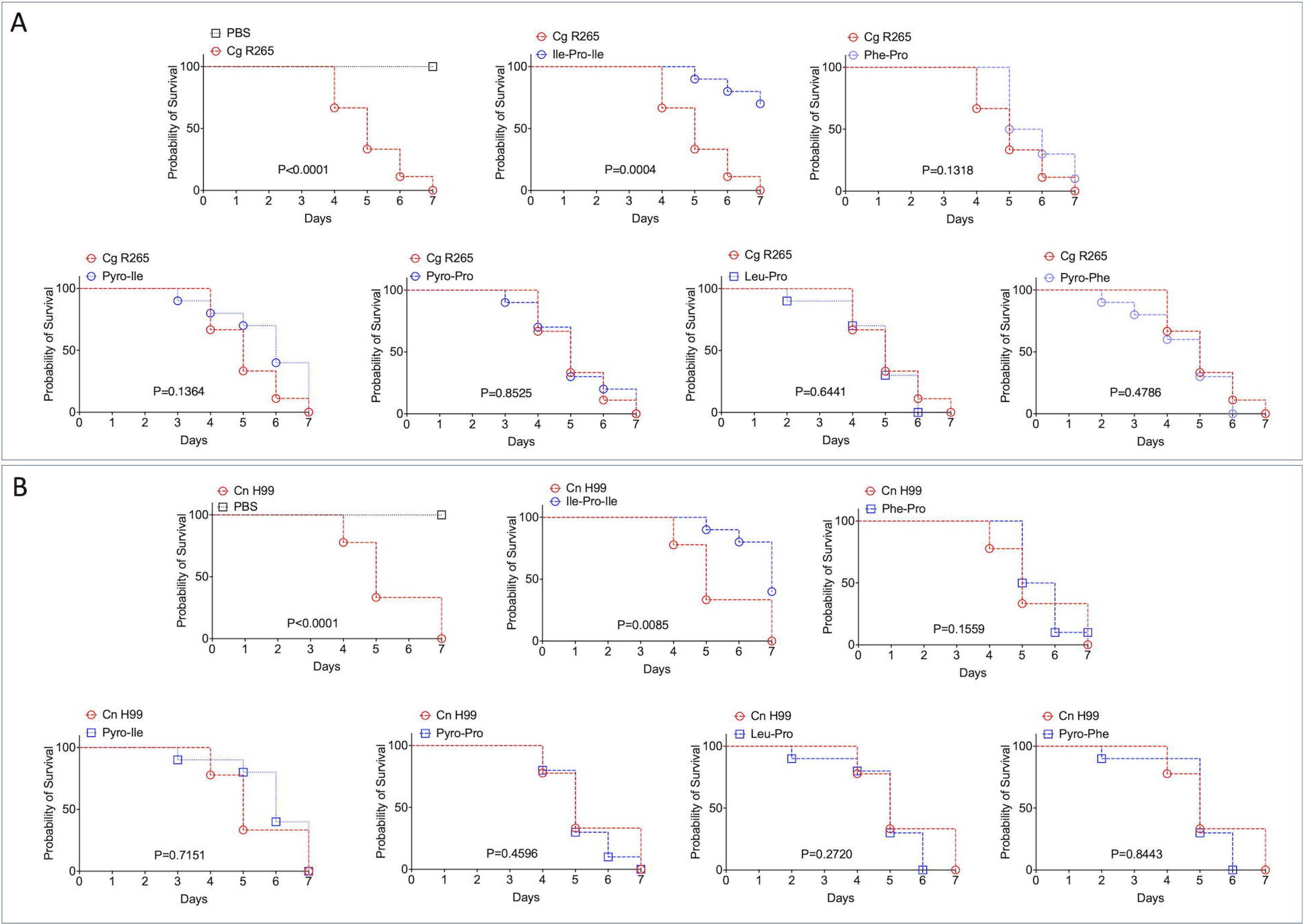
Effects of the EV peptides (10 μg/ml) on the survival of *G. mellonella* lethally infected with *C. gattii* R265 (Cg; A) or *C. neoformans* H99 (Cn; B). A. Ile-Pro-Ile was the only peptide prolonging the survival of *G. mellonella*. The other peptides did not interfere with the host’s survival. The experiment illustrated in A was repeated using *C. neoformans* H99 instead of *C. gattii* R265, producing similar results.

Phe-Pro, Pyro-Iso, Pyr-Pro, Leu-Pro and Pyro-Phe did not have any effect on the survival curves. In contrast, the tripeptide Ile-Pro-Ile significantly improved the survival of *G. mellonella*. We repeated this experiment using *C. neoformans* instead of *C. gattii* and obtained similar results (Figure 5B). On the basis of these results, we selected Ile-Pro-Ile for tests at lower concentrations (1, 0.5 and 0.1 μg/ml) in the *C. gattii* infection model. Once again, the peptide was highly efficient in prolonging the survival of lethally infected *G. mellonella* in a dose-dependent fashion (Figure 6).

**Figure 6.**
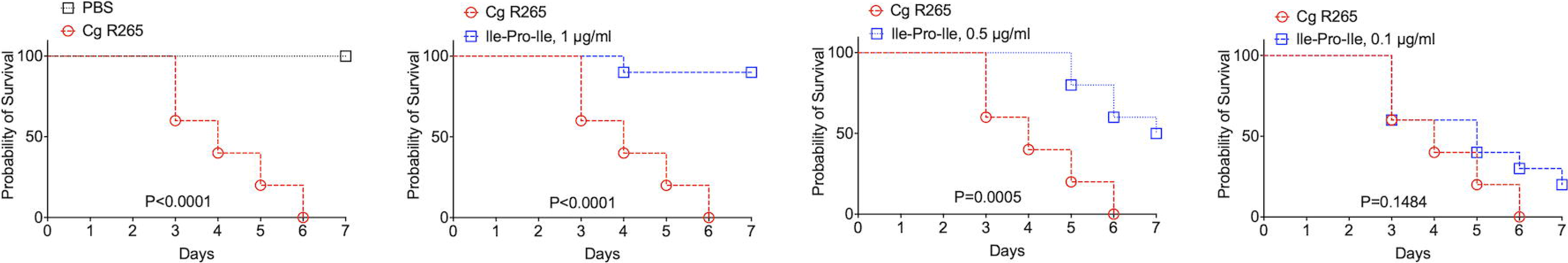
Dose-dependent protection of *G. mellonella* against *C. gattii* (Cg) induced by Ile-Pro-Ile. Survival of *G. mellonella* after injection with PBS (control) or with *C. gattii* yeast cells (left panel) is shown, in addition to the comparative survival curves of *G. mellonella* after injection with *C. gattii* alone (red curves) or with the fungus in the presence of variable concentrations of Ile-Pro-Ile.

## Discussion

The knowledge of the functions of fungal EVs has continuously increased in the recent years (7), but the biological roles of low mass structures exported in EVs are unknown. Small molecules secreted by *Cryptococcus* are immunologically active and affect IL-1β inflammasome-dependent secretion (24), but their association with EVs has not been established. In our study, we aimed at proving the concept that biologically active small molecules are exported in cryptococcal EVs. This idea culminated with the characterization for the first time of a fungal EV molecule inducing protection against pathogenic fungi.

Fungal EVs were demonstrated to mediate intercellular communication (4), prion transmission (25), biofilm formation associated with antifungal resistance (26), immunological responses *in vitro* (23,27–30), and protection of different hosts against lethal challenges with fungal pathogens (12,21–23). In any of these examples, these biological effects attributed to the EVs were correlated with the identification of the bioactive vesicular molecules. The only known exception was the protection of *G. mellonella* induced by cryptococcal EVs enriched with sterol glycosides and capsular polysaccharides (22). However, it is important to mention that the EVs in this study were produced by genetically engineered cells and, therefore, did not correspond to native vesicles. It remained also unknown if other molecules influenced the protective effects, since compositional studies have not been performed.

The identification of bioactive EV molecules is challenging in multiple aspects. The compositional analysis of fungal EVs in different models include a formidable variability in culture conditions, since each of the fungal pathogens tested so far manifest growth particularities. In this scenario, biomarkers of fungal EVs are still not known, although it has been suggested that mannoproteins and claudin-like Sur7 family proteins are important components of vesicles produced by *C. neoformans* and *C. albicans*, respectively (12,31). The knowledge of small molecules mediating important biological activities in fungal EVs is even more limited. In *H. capsulatum*, carbohydrate metabolites were abundantly detected in EVs, in addition to L-ornithine and ethanolamine, among other small molecules (10). In the plant pathogen *P. digitatum*, EVs were characterized as the carriers of tryptoquialanine A, a toxin that inhibited the germination of orange seeds (11). So far, tryptoquialanine A is the only low mass component of fungal EVs with a reported function. In this model, the mycotoxin fungisporin was also detected (11), but its function in fungal EVs remains to be determined. Together, these findings illustrate the need for an improved knowledge of the composition and functions of EV metabolites in fungi.

The isolation of cryptococcal EVs from solid medium is much more efficient than the similar protocols using liquid cultures (5). RNA and proteins in cryptococcal EVs obtained in liquid cultures were characterized in early studies (8,32), but their distribution in EVs obtained from solid medium was only recently described in *C. neoformans* (12). Other molecules remained unknown, and the metabolite composition of cryptococcal EVs has not been investigated so far. In our study, we initially aimed at understanding what are the low molecular weight components exported by *C. gattii* in solid medium. We identified small molecules of different chemical natures as putative components of cryptococcal EVs, but their functions remain widely unknown. However, our chemical and biological methods for structural validation revealed that one tripeptide was capable to protect *G. mellonella* against lethal challenges with *C. gattii* or *C. neoformans*. The mechanisms by which the peptides induced protection against cryptococcal infection remain unknown, but the immune response of *G. mellonella* is innate and relies on the activity of hemocytes in combination with antimicrobial peptides and lytic enzymes, among others (33).

Accordingly, immunity to *Cryptococcus* relies on innate immune cells coordinating adaptive responses to stimulate fungal killing (34). Therefore, we hypothesize that the tripeptide identified in our study is an inducer of innate responses, which have a key general role in the control of fungal infections (35).

The peptide inducing protection against *Cryptococcus* in *G. mellonella* was demonstrated to have important biological activities in other models. Ile-Pro-Ile, also known as diprotin A, is an inhibitor of dipeptidyl peptidase 4, an enzyme participating in insulin metabolism (36) and chemotaxis of murine embryonic stem cells towards stromal cell-derived factor-1 (37). Its role in fungal physiology and/or pathogenesis are still unknown. Noteworthy, our study did not elucidate any physiological or pathogenic functions. Instead, we present a proof of concept that fungal EVs are the vehicles for exporting biologically active molecules of low molecular mass that may be involved in immunological and/or pathogenic mechanisms. Since fungal EVs have been consistently proposed as vaccine candidates in different models, the potential of these findings can be substantial.

## Methods

### Preparation of EVs

The EV-producing isolate used in this study was the standard strain R265 of *C. gattii*. Of note, the R265 strain has been recently reclassified as *C. deuterogattii* (38). In this study, we kept its classification as *C. gattii*, as largely employed in the *Cryptococcus* literature. EV isolation was based on the protocol that we have recently established for *C. gattii* and other fungal species (5). Briefly, One colony of *C. gattii* R265 cultivated in solid Sabouraud’s medium was inoculated into yeast extract-peptone-dextrose (YPD) medium (5 ml) and cultivated for 1 day at 30°C with shaking. The cell density was adjusted to of 3.5× 10^7^cells/ml in YPD. From this suspension, aliquots of 300 μl were taken for inoculation in YPD agar plates, which were cultivated for 1 day at 30°C. The cells were recovered from the plates with an inoculation loop and transferred to a single centrifuge tube containing 30 ml of PBS filtered through 0.22-μm-pore membranes. The cells were then removed by centrifugation (5,000 × g for 15 min at 4°C), and the supernatants were centrifuged again (15,000 × g for 15 min at 4°C) to remove debris. The resulting supernatants were filtered through 0.45-μm-pore syringe filters and again centrifuged (100,000 × g, 1 h at 4°C). Supernatants were discarded and pellets suspended in 300 μl of sterile PBS. To avoid the characterization of medium components as EV molecules, mock (control) samples were similarly prepared using sterile plates containing YPD. Four petri dishes were used for each EV isolation, and EV isolation was performed independently three times. In all samples, the properties of EVs and their concentration was monitored by nanoparticle tracking analysis (NTA) as described by our group (5). The samples prepared for mass spectrometry analyses had the typical properties of *C. gattii* EVs (data not shown), and were in the range of 4 to 6 × 10^10^ EVs within the triplicate set.

### Mass spectrometry analyses

*C. gattii* EVs were vacuum dried and extracted with 1 ml of methanol during 1 h in an ultrasonic bath. The extracts were filtered (0.22 μm), dried under a gentle N2 flux and stored at −20 °C. EV extracts were resuspended in 200 μl of MeOH and transferred into glass vials. Ultra high-performance liquid chromatography-mass spectrometry (UHPLC-MS) analyses were performed using a Thermo Scientific QExactive^®^ hybrid Quadrupole-Orbitrap mass spectrometer with the following parameters: electrospray ionization in positive mode, capillary voltage at 3.5 kV; capillary temperature at 300 °C; S-lens of 50 V and *m/z* range of 100.00-1500.00. Tandem Mass spectrometry (MS/MS) was performed using normalized collision energy (NCE) of 20, 30 and 40 eV; maximum 5 precursors per cycle were selected. Stationary phase was a Waters ACQUITY UPLC^®^ BEH C18 1.7 μm (2.1 mm × 50 mm) column.

Mobile phases were 0.1 % (v/v) formic acid in water (A) and acetonitrile (B). Eluent profile (A:B) 0-10 min, gradient from 95:5 up to 2:98; held for 5 min; 15-16.2 min gradient up to 95:5; held for 3.8 min. Flow rate was 0.2 mL min^−1^. Injection volume was 3 μL. UHPLC-MS operation and spectra analyses were performed using Xcalibur software (version 3.0.63).

### Molecular Network

A molecular network was created using the online workflow (https://ccms-ucsd.github.io/GNPSDocumentation/) on the GNPS website (http://gnps.ucsd.edu). The data was filtered by removing all MS/MS fragment ions within +/− 17 Da of the precursor m/z. MS/MS spectra were window filtered by choosing only the top 6 fragment ions in the +/− 50Da window throughout the spectrum. The precursor ion mass tolerance was set to 0.02 Da and a MS/MS fragment ion tolerance of 0.02 Da. A network was then created where edges were filtered to have a cosine score above 0.5 and more than 5 matched peaks. Further, edges between two nodes were kept in the network if and only if each of the nodes appeared in each other’s respective top 10 most similar nodes. Finally, the maximum size of a molecular family was set to 100, and the lowest scoring edges were removed from molecular families until the molecular family size was below this threshold. The spectra in the network were then searched against GNPS’ spectral libraries. The library spectra were filtered in the same manner as the input data. All matches kept between network spectra and library spectra were required to have a score above 0.5 and at least 5 matched peaks (14).

### Peptides

The peptides selected for biological tests were synthesized by GenOne Biotechnologies (https://www.genone.com.br, Rio de Janeiro, Brazil). Purity and structural properties of each peptide were confirmed by high-performance liquid chromatography coupled to mass spectrometry. All peptides were water-soluble and had their purity at the 95% range.

### *Galleria mellonella* infection model

Groups of 10 larvae (250 – 350 mg) were infected with 10 μl of sterile PBS containing 10^6^ cells of *C. gattii* or *C. neoformans* into the last left proleg using a Hamilton micro-syringe. Control systems were injected with PBS alone. Fungal inoculation was followed by injection of 10 μl PBS solutions containing Ile-Pro-Ile, Phe-Pro, Pyro-Ile, Pyro-Pro, Leu-Pro, and Pyro-Phe at 10 μg/ml. Due to its promising effects, Ile-Pro-Ile was also tested at 1, 0.5, and 0.1 μl/ml in a *C. gattii* model of infection. Injected larvae were placed in sterile Petri dishes and incubated at 37°C. The survival was monitored daily in a period of seven days. Larvae were considered dead if they did not respond to physical stimulus. Statistical analysis was performed using the Graphpad Prism software, version 8.0.

## Acknowledgements

M.L.R. is currently on leave from the position of associate professor at the Microbiology Institute of the Federal University of Rio de Janeiro, Brazil. M.L.R. is supported by grants from the Brazilian Ministry of Health (grant 440015/2018-9), Conselho Nacional de Desenvolvimento Científico e Tecnológico (CNPq; grants 405520/2018-2 and 301304/2017-3), and Fiocruz (grants PROEP-ICC 442186/2019-3, VPPCB-007-FIO-18, and VPPIS-001-FIO18). JHC and FCGR were financed in part by scholarships from the Coordenação de Aperfeiçoamento de Pessoal de Nível Superior (CAPES, Brazil, Finance Code 001). MLR also acknowledges support from the Instituto Nacional de Ciência e Tecnologia de Inovação em Doenças de Populações Negligenciadas (INCT-IDPN). The funders had no role in the decision to publish, or preparation of the manuscript.

## Conflict of interest statement

The authors report no conflict of interest.

## Notes

### Competing Interest Statement

The authors have declared no competing interest.

## References

1. Kwon-Chung KJ, Fraser JA, Doering TÁL, Wang ZA, Janbon G, Idnurm A, Bahn YS. Cryptococcus neoformans and Cryptococcus gattii, the etiologic agents of cryptococcosis. Cold Spring Harb Perspect Med (2015) doi:10.1101/cshperspect.a019760

2. Fernando Silva Rocha D, Cruz KS, da Silva Santos CS, Stephanny Fernandes Menescal L, da Silva Neto JR, Pinheiro SB, Silva LM, Trilles L, Vicente Braga de Souza J. MLST reveals a clonal population structure for Cryptococcus neoformans molecular type VNI isolates from clinical sources in Amazonas, Northern-Brazil. PLoS One (2018) doi:10.1371/journal.pone.0197841

3. Hagen F, Ceresini PC, Polacheck I, Ma H, van Nieuwerburgh F, Gabaldón T, Kagan S, Pursall ER, Hoogveld HL, van Iersel LJJ, et al. Ancient Dispersal of the Human Fungal Pathogen Cryptococcus gattii from the Amazon Rainforest. PLoS One (2013) doi:10.1371/journal.pone.0071148

4. Bielska E, Sisquella MA, Aldeieg M, Birch C, O’Donoghue EJ, May RC. Pathogen-derived extracellular vesicles mediate virulence in the fatal human pathogen Cryptococcus gattii. Nat Commun (2018) 9: doi:10.1038/s41467-018-03991-6

5. Reis FCG, Borges BS, Jozefowicz LJ, Sena BAG, Garcia AWA, Medeiros LC, Martins ST, Honorato L, Schrank A, Vainstein MH, et al. A novel protocol for the isolation of fungal extracellular vesicles reveals the participation of a putative scramblase in polysaccharide export and capsule construction in Cryptococcus gattii. mSphere (2019) 4:e00080–19. doi:10.1128/mSphere.00080-19

6. Rodrigues ML, Nimrichter L, Oliveira DL, Frases S, Miranda K, Zaragoza O, Alvarez M, Nakouzi A, Feldmesser M, Casadevall A. Vesicular polysaccharide export in Cryptococcus neoformans is a eukaryotic solution to the problem of fungal trans-cell wall transport. Eukaryot Cell (2007) 6: doi:10.1128/EC.00318-06

7. Rizzo J, Rodrigues ML, Janbon G. Extracellular Vesicles in Fungi: Past, Present, and Future Perspectives. Front Cell Infect Microbiol (2020) 10: doi:10.3389/fcimb.2020.00346

8. Rodrigues ML, Nakayasu ES, Almeida IC, Nimrichter L. The impact of proteomics on the understanding of functions and biogenesis of fungal extracellular vesicles. J Proteomics (2014)97:177–186. doi:10.1016/j.jprot.2013.04.001

9. de Toledo Martins S, Szwarc P, Goldenberg S, Alves LR. Extracellular Vesicles in Fungi: Composition and Functions. Curr Top Microbiol Immunol (2019) 422:45–59. doi:10.1007/82_2018_141

10. Cleare LG, Zamith D, Heyman HM, Couvillion SP, Nimrichter L, Rodrigues ML, Nakayasu ES, Nosanchuk JD. Media matters! Alterations in the loading and release of Histoplasma capsulatum extracellular vesicles in response to different nutritional milieus. Cell Microbiol (2020) doi:10.1111/cmi.13217

11. Costa JH, Bazioli JM, Barbosa LD, dos Santos Júnior PLT, Reis FCG, Klimeck T, Crnkovic CM, Berlinck RGS, Sussulini A, Rodrigues ML, et al. Phytotoxic tryptoquialanines produced *in vivo* by *Penicillium digitatum* are exported in extracellular vesicles. bioRxiv (2020)2020.12.03.411132. doi:10.1101/2020.12.03.411132

12. Rizzo juliana, Wong SSW, Gazi AD, Moyrand F, Chaze T, Commere P-H, Matondo M, Novault S, Pehau-Arnaudet G, Reis F, et al. New insights into Cryptococcus extracellular vesicles suggest a new structural model and an antifungal vaccine strategy. bioRxiv (2020)2020.08.17.253716. doi:10.1101/2020.08.17.253716

13. Rodrigues ML, Nakayasu ES, Oliveira DL, Nimrichter L, Nosanchuk JD, Almeida IC, Casadevall A. Extracellular vesicles produced by Cryptococcus neoformans contain protein components associated with virulence. Eukaryot Cell (2008) 7: doi:10.1128/EC.00370-07

14. Wang M, Carver JJ, Phelan V V., Sanchez LM, Garg N, Peng Y, Nguyen DD, Watrous J, Kapono CA, Luzzatto-Knaan T, et al. Sharing and community curation of mass spectrometry data with Global Natural Products Social Molecular Networking. Nat Biotechnol (2016) doi:10.1038/nbt.3597

15. Krug D, Müller R. Secondary metabolomics: The impact of mass spectrometry-based approaches on the discovery and characterization of microbial natural products. Nat Prod Rep (2014) doi:10.1039/c3np70127a

16. Purves K, Macintyre L, Brennan D, Hreggviðsson G, Kuttner E, Ásgeirsdóttir ME, Young LC, Green DH, Edrada-Ebel R, Duncan KR. Using molecular networking for microbial secondary metabolite bioprospecting. Metabolites (2016) doi:10.3390/metabo6010002

17. Watrous J, Roach P, Alexandrov T, Heath BS, Yang JY, Kersten RD, Van Der Voort M, Pogliano K, Gross H, Raaijmakers JM, et al. Mass spectral molecular networking of living microbial colonies. Proc Natl Acad Sci U S A (2012) doi:10.1073/pnas.1203689109

18. Nguyen DD, Wu CH, Moree WJ, Lamsa A, Medema MH, Zhao X, Gavilan RG, Aparicio M, Atencio L, Jackson C, et al. MS/MS networking guided analysis of molecule and gene cluster families. Proc Natl Acad Sci U S A (2013) doi:10.1073/pnas.1303471110

19. Zhang P, Chan W, Ang IL, Wei R, Lam MMT, Lei KMK, Poon TCW. Revisiting Fragmentation Reactions of Protonated α-Amino Acids by High-Resolution Electrospray Ionization Tandem Mass Spectrometry with Collision-Induced Dissociation. Sci Rep (2019) doi:10.1038/s41598-019-42777-8

20. Kombrink A, Tayyrov A, Essig A, Stöckli M, Micheller S, Hintze J, van Heuvel Y, Dürig N, Lin C wei, Kallio PT, et al. Induction of antibacterial proteins and peptides in the coprophilous mushroom Coprinopsis cinerea in response to bacteria. ISME J (2019) doi:10.1038/s41396-018-0293-8

21. Vargas G, Honorato L, Guimarães AJ, Rodrigues ML, Reis FCG, Vale AM, Ray A, Nosanchuk JD, Nimrichter L. Protective effect of fungal extracellular vesicles against murine candidiasis. Cell Microbiol (2020) doi:10.1111/cmi.13238

22. Ana Caroline Colombo, Rella A, Normile T, Joffe LS, Tavares PM, Glauber GR, Frases S, Orner EP, Farnoud AM, Fries BC, et al. Cryptococcus neoformans glucuronoxylomannan and sterylglucoside are required for host protection in an animal vaccination model. MBio (2019) doi:10.1128/mBio.02909-18

23. Vargas G, Rocha JDB, Oliveira DL, Albuquerque PC, Frases S, Santos SS, Nosanchuk JD, Gomes AMO, Medeiros LCAS, Miranda K, et al. Compositional and immunobiological analyses of extracellular vesicles released by Candida albicans. Cell Microbiol (2015) 17:389–407. doi:10.1111/cmi.12374

24. Bürgel PH, Marina CL, Saavedra PHV, Albuquerque P, Holanda PH, de Araújo Castro R, Heyman H, Coelho C, Cordero RJB, Casadevall A, et al. Cryptococcus neoformans secretes small molecules that inhibit IL-1β inflammasome-dependent secretion. bioRxiv (2019) doi:10.1101/554048

25. Kabani M, Melki R. Sup35p in its soluble and prion states is packaged inside extracellular vesicles. MBio (2015) 6: doi:10.1128/mBio.01017-15

26. Zarnowski R, Sanchez H, Covelli AS, Dominguez E, Jaromin A, Bernhardt J, Mitchell KF, Heiss C, Azadi P, Mitchell A, et al. Candida albicans biofilm–induced vesicles confer drug resistance through matrix biogenesis. PLOS Biol (2018) doi:10.1371/journal.pbio.2006872

27. Oliveira DL, Freire-de-Lima CG, Nosanchuk JD, Casadevall A, Rodrigues ML, Nimrichter L. Extracellular vesicles from Cryptococcus neoformans modulate macrophage functions. Infect Immun (2010) doi:10.1128/IAI.01171-09

28. Da Silva TA, Roque-Barreira MC, Casadevall A, Almeida F. Extracellular vesicles from Paracoccidioides brasiliensis induced M1 polarization in vitro. Sci Rep (2016) doi:10.1038/srep35867

29. Bitencourt TA, Rezende CP, Quaresemin NR, Moreno P, Hatanaka O, Rossi A, Martinez-Rossi NM, Almeida F. Extracellular vesicles from the dermatophyte trichophyton interdigitalemodulate macrophage and keratinocyte functions. Front Immunol (2018) doi:10.3389/fimmu.2018.02343

30. Almeida F, Wolf JM, Da Silva TA, Deleon-Rodriguez CM, Rezende CP, Pessoni AM, Fernandes FF, Silva-Rocha R, Martinez R, Rodrigues ML, et al. Galectin-3 impacts Cryptococcus neoformans infection through direct antifungal effects. Nat Commun (2017) doi:10.1038/s41467-017-02126-7

31. Dawson CS, Garcia-Ceron D, Rajapaksha H, Faou P, Bleackley MR, Anderson MA. Protein markers for Candida albicans EVs include claudin-like Sur7 family proteins. J Extracell Vesicles (2020) doi:10.1080/20013078.2020.1750810

32. Da Silva RP, Puccia R, Rodrigues ML, Oliveira DL, Joffe LS, César G V., Nimrichter L, Goldenberg S, Alves LR. Extracellular vesicle-mediated export of fungal RNA. Sci Rep (2015) 5:7763. doi:10.1038/srep07763

33. Trevijano-Contador N, Zaragoza O. Immune response of Galleria mellonella against human fungal pathogens. J Fungi (2019) doi:10.3390/jof5010003

34. Gibson JF, Johnston SA. Immunity to Cryptococcus neoformans and C. gattii during cryptococcosis. Fungal Genet Biol (2015) doi:10.1016/j.fgb.2014.11.006

35. Shoham S, Levitz SM. The immune response to fungal infections. Br J Haematol (2005) doi:10.1111/j.1365-2141.2005.05397.x

36. Kieffer TJ, Mc Intosh CHS, Pederson RA. Degradation of glucose-dependent insulinotropic polypeptide and truncated glucagon-like peptide 1 in vitro and in vivo by dipeptidyl peptidase iv. Endocrinology (1995) doi:10.1210/endo.136.8.7628397

37. Guo Y, Hangoc G, Bian H, Pelus LM, Broxmeyer HE. SDF-1/CXCL12 Enhances Survival and Chemotaxis of Murine Embryonic Stem Cells and Production of Primitive and Definitive Hematopoietic Progenitor Cells. Stem Cells (2005) doi:10.1634/stemcells.2005-0085

38. Hagen F, Khayhan K, Theelen B, Kolecka A, Polacheck I, Sionov E, Falk R, Parnmen S, Lumbsch HT, Boekhout T. Recognition of seven species in the Cryptococcus gattii/Cryptococcus neoformans species complex. Fungal Genet Biol (2015) doi:10.1016/j.fgb.2015.02.009

