## Supplemental Figures 1-13 for "Small molecule analysis of extracellular vesicles produced by *Cryptococcus gattii*: identification of a tripeptide controlling cryptococcal infection in an invertebrate host model"

**Supplemental material**

**Supplemental Figures 1-13**. Metabolite identification through hits in the GNPS database. MS data are shown for Ile-Pro-Ile (Figure S1), Phe-Pro (Figure S2), pyro-Glu-Ile (Figure S3), pyro-Glu-Pro (Figure S4), Leu-Pro (Figure S5), pyro-Glu-Phe (Figure S6), Val-Leu-Pro-Val-Pro (Figure S7), cyclo(Trp-Pro) (Figure S8), cyclo(Tyr-Pro) (Figure S9), tryptophan (Figure S10), asperphenamate (Figure S11), riboflavin (Figure S12), and pantothenic acid (Figure S13).

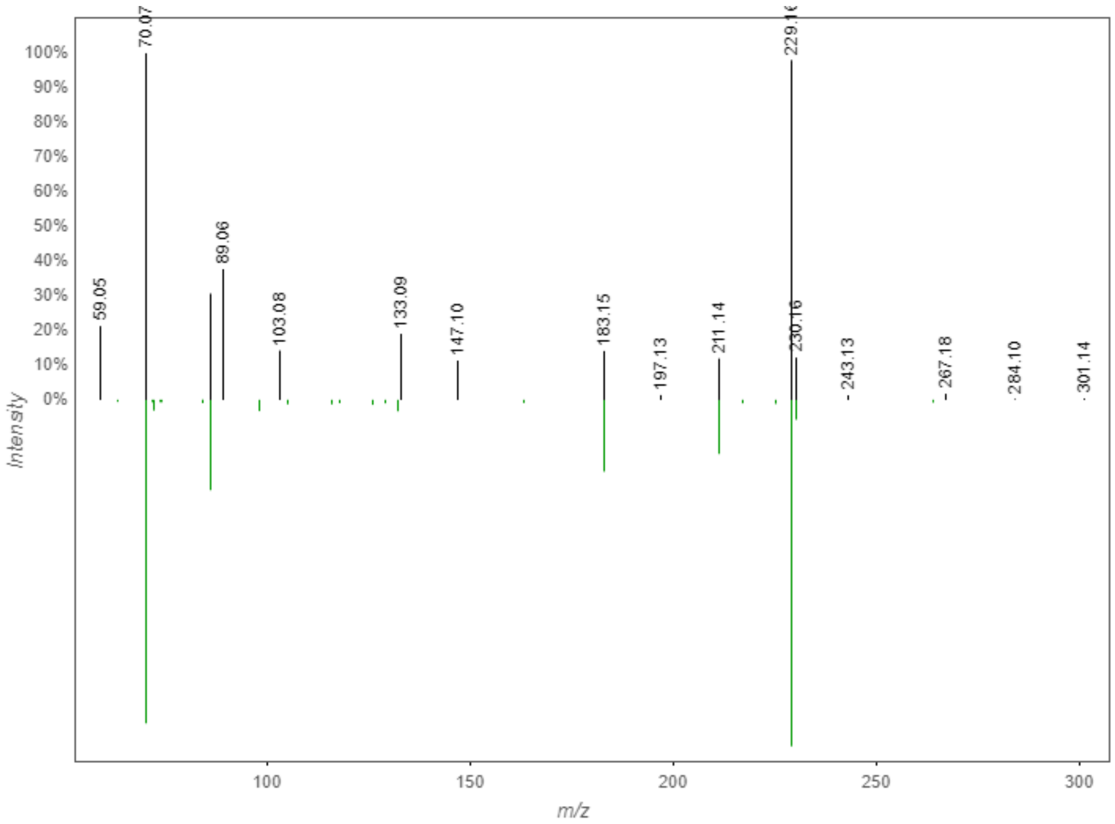


Figure S1. GNPS hit/MS data for Ile-Pro-Ile.

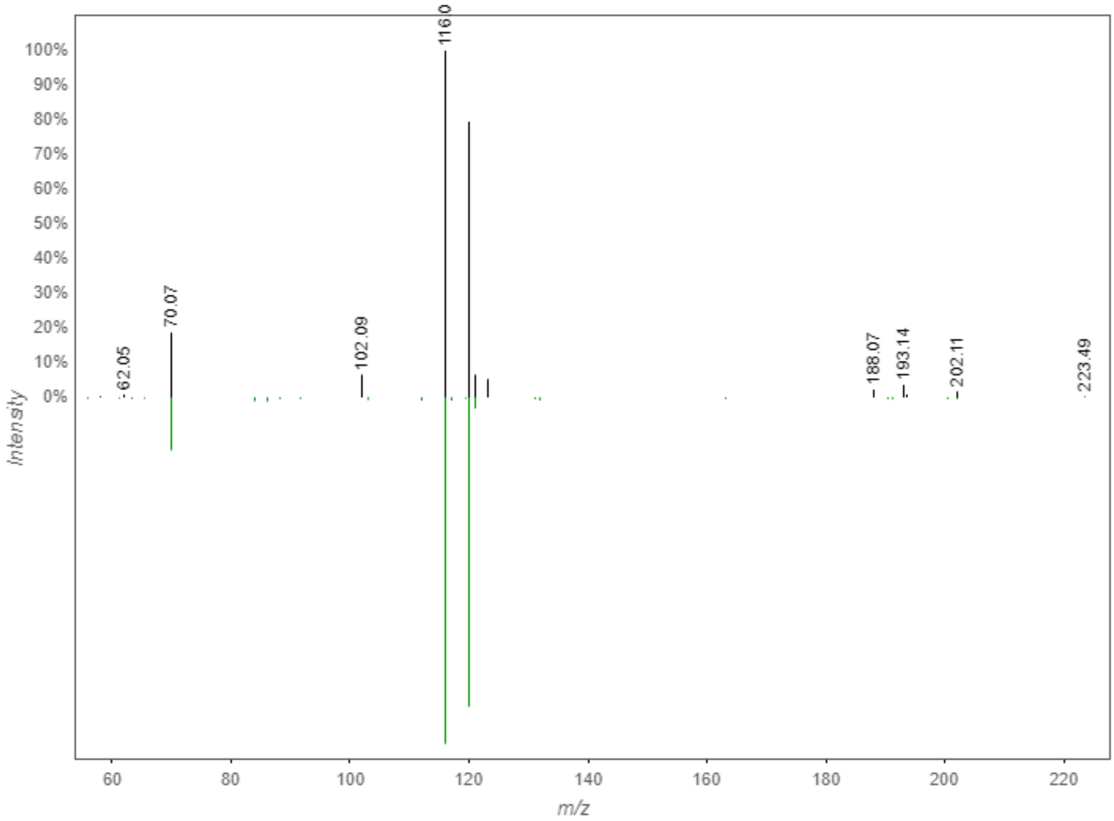


Figure S2. GNPS hit/MS data for Phe-Pro.

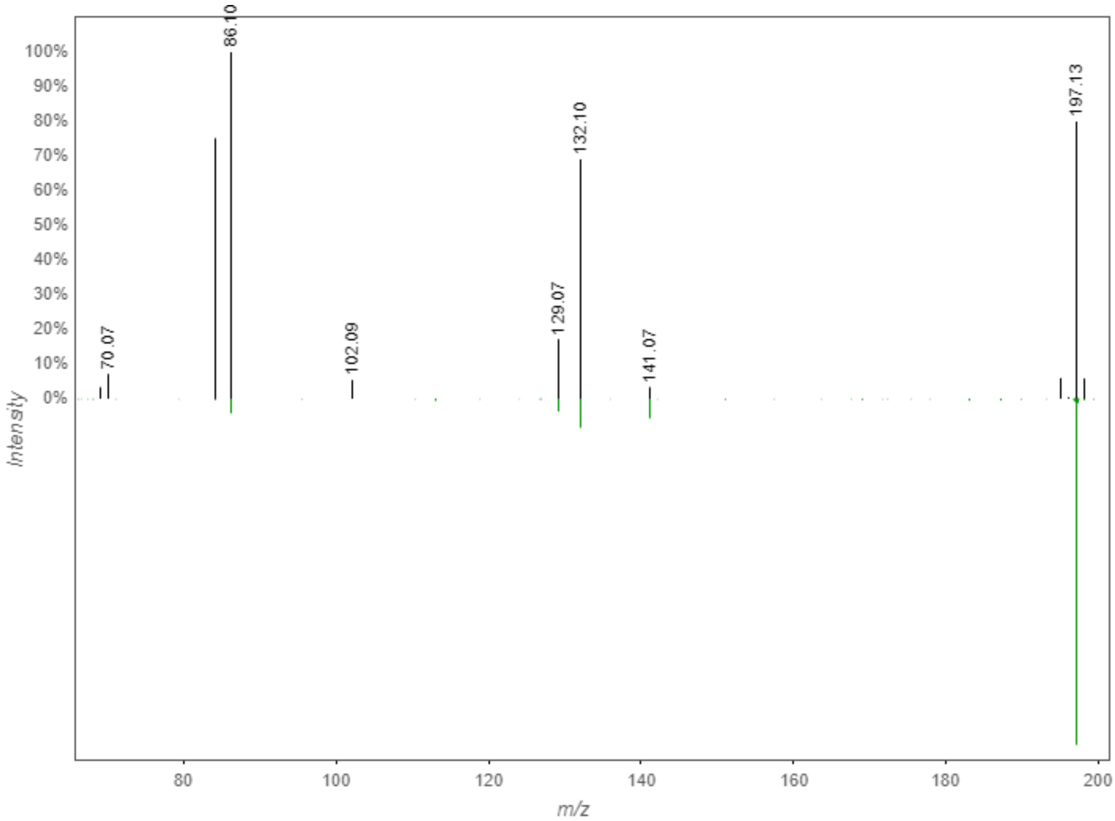

Figure S3. GNPS hit/MS data for pyro-Glu-Ile.

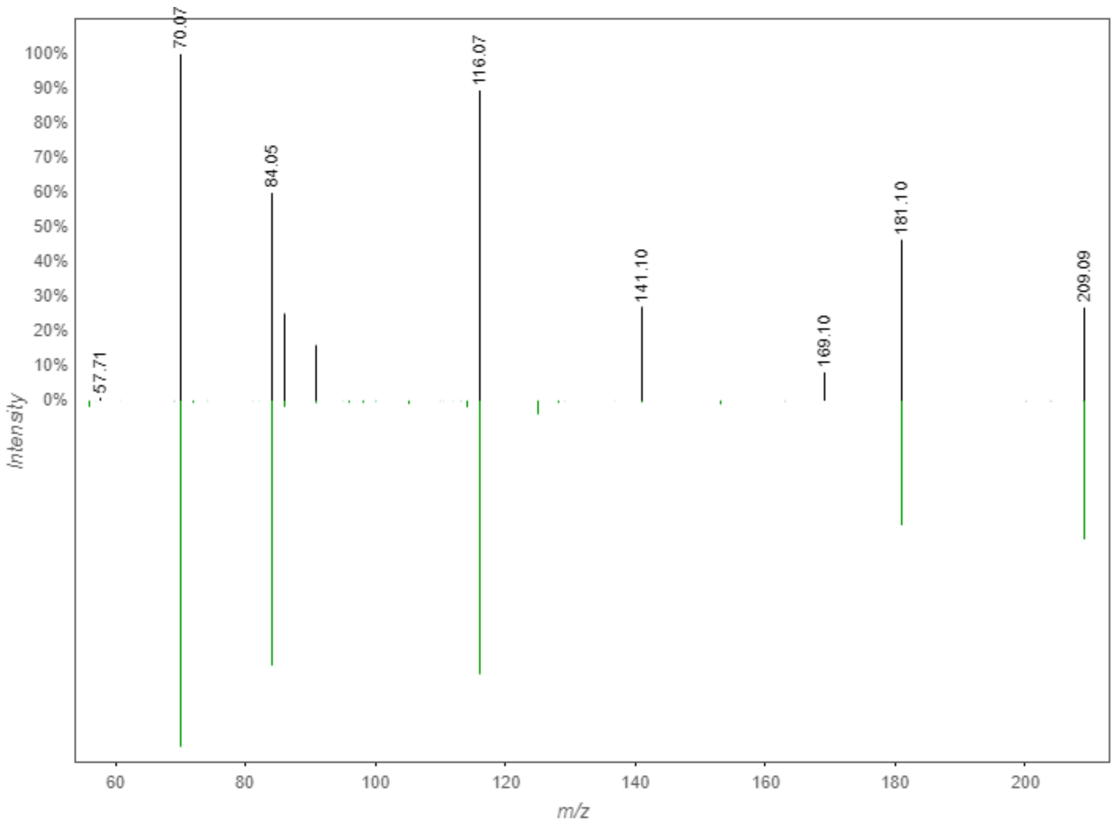

Figure S4. GNPS hit/MS data for pyro-Glu-Pro.

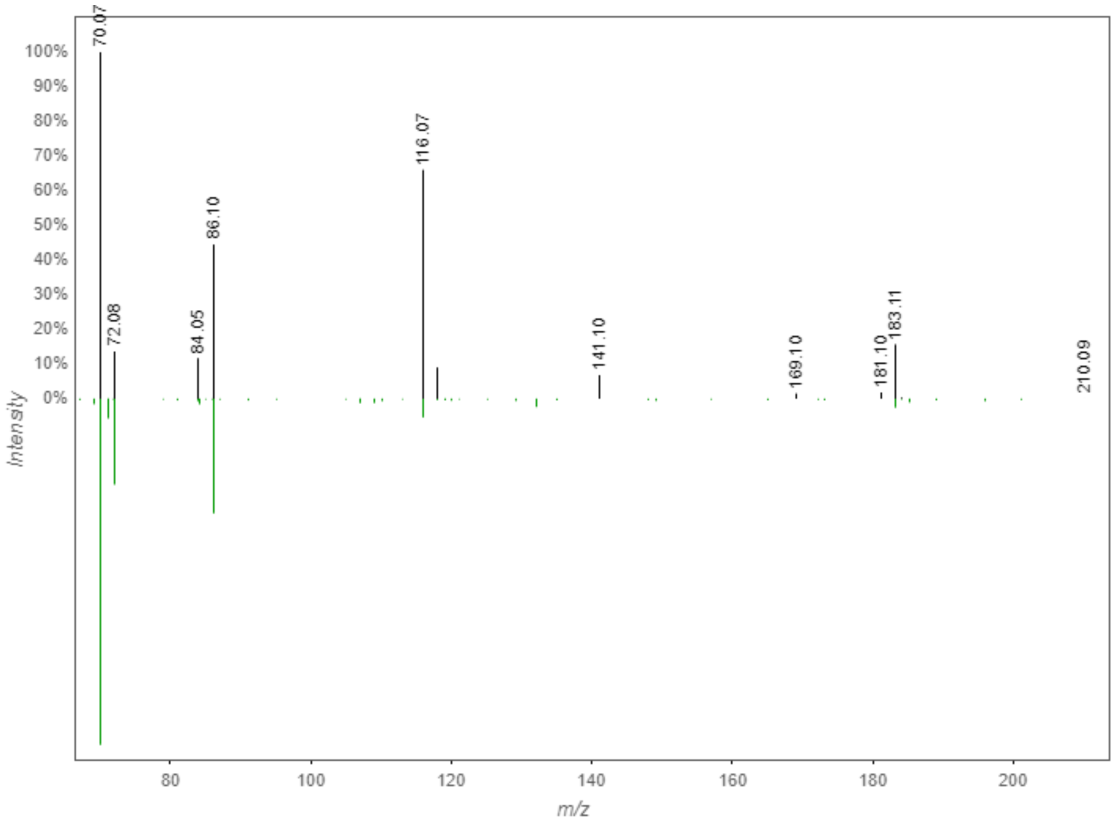


Figure S5. GNPS hit/MS data for Leu-Pro.

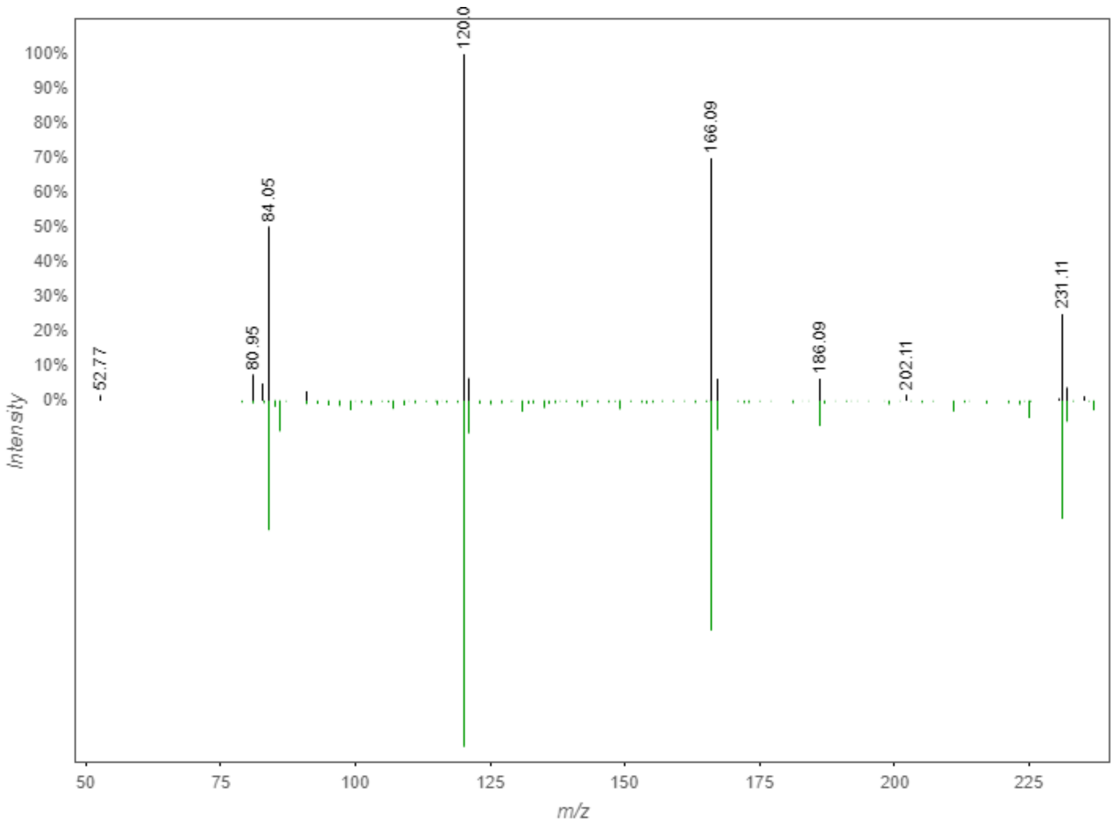

Figure S6. GNPS hit/MS data for pyro-Glu-Phe.

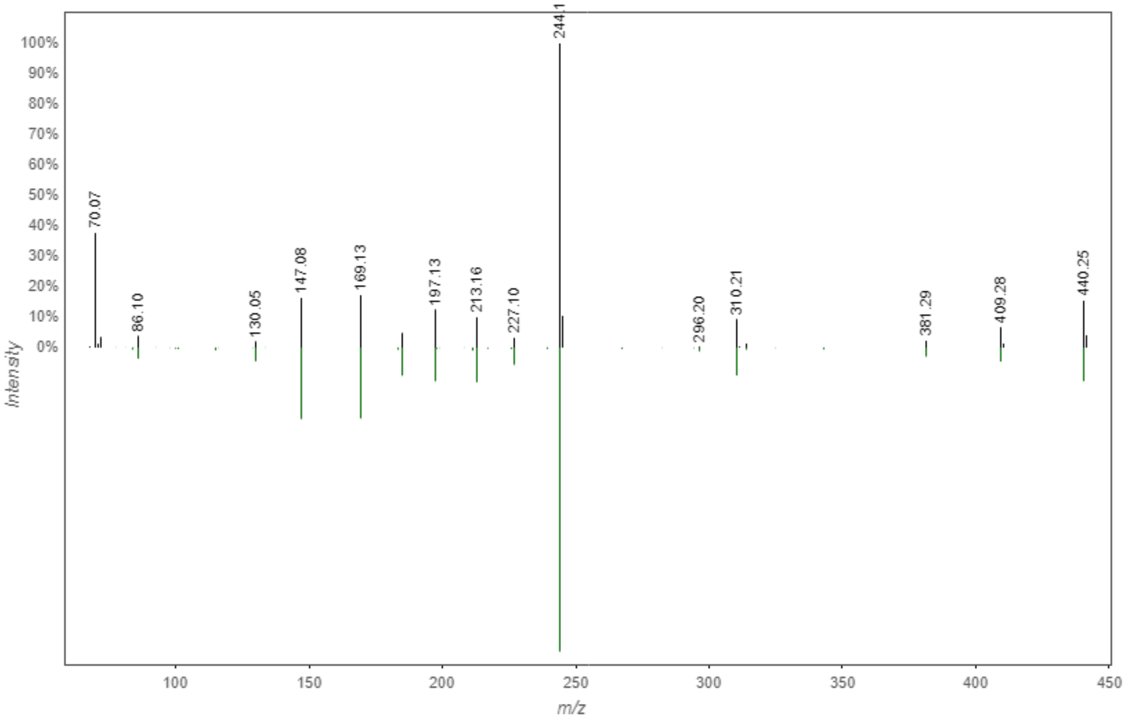

Figure S7. GNPS hit/MS data for Val-Leu-Pro-Val-Pro.

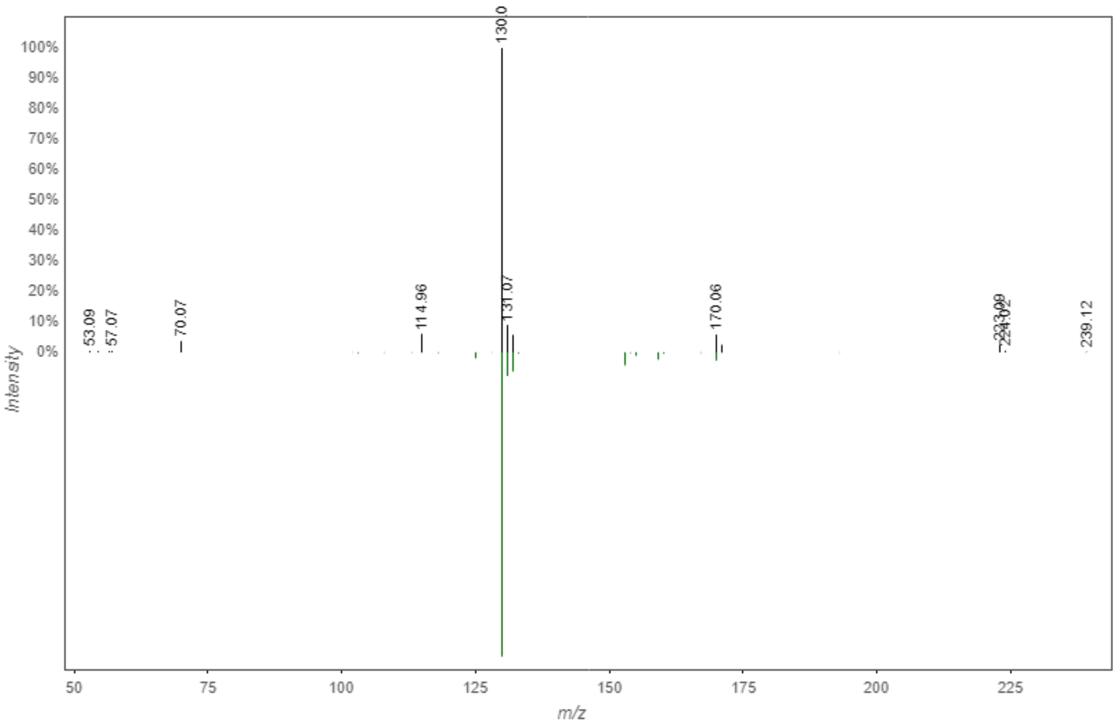


Figure S8. GNPS hit/MS data for cyclo(Trp-Pro).

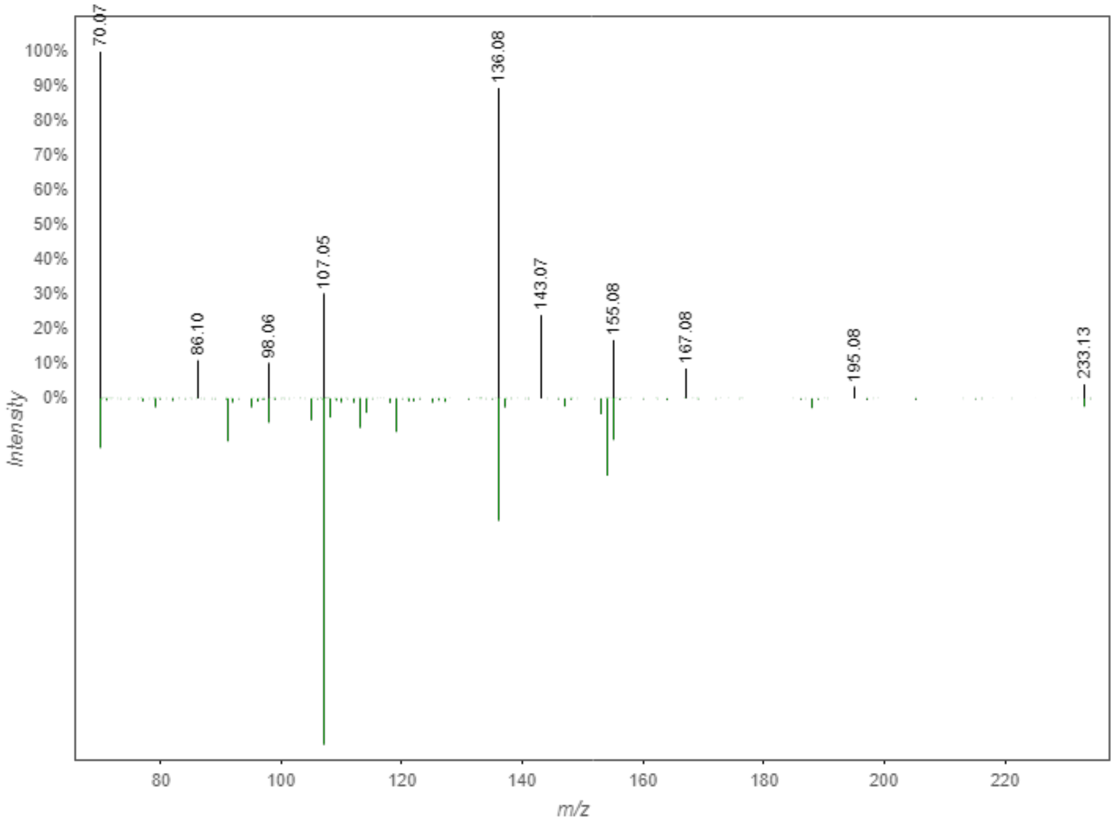

Figure S9. GNPS hit/MS data for cyclo(Tyr-Pro).

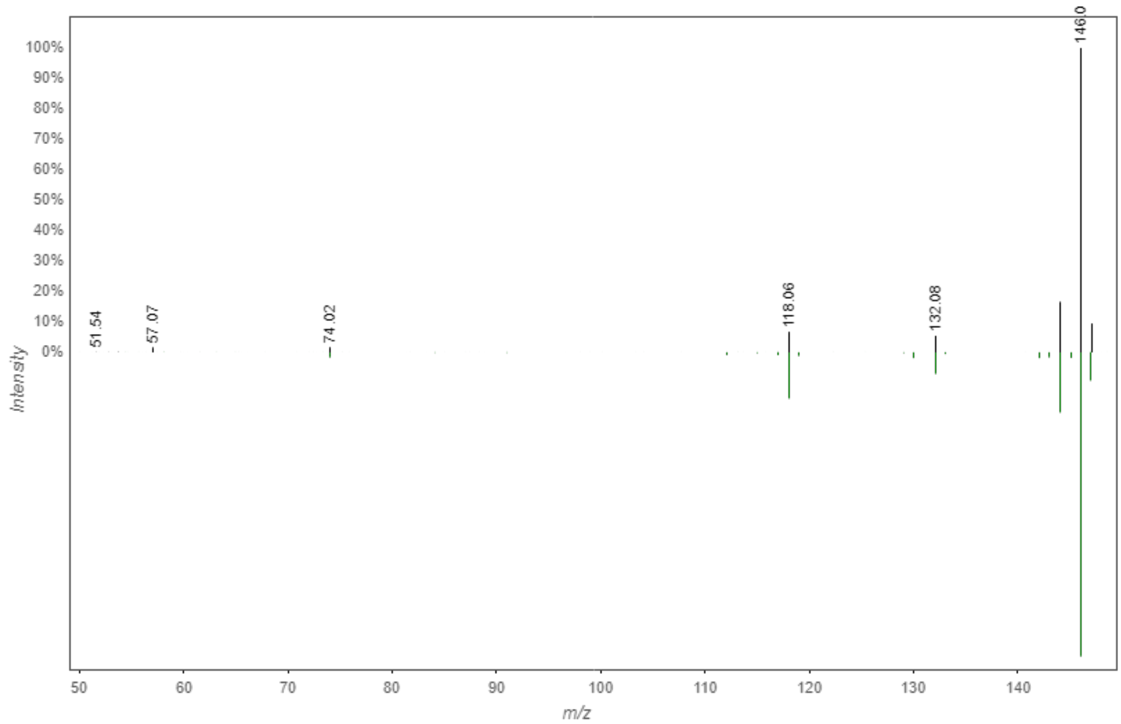

Figure S10. GNPS hit/MS data for tryptophan.

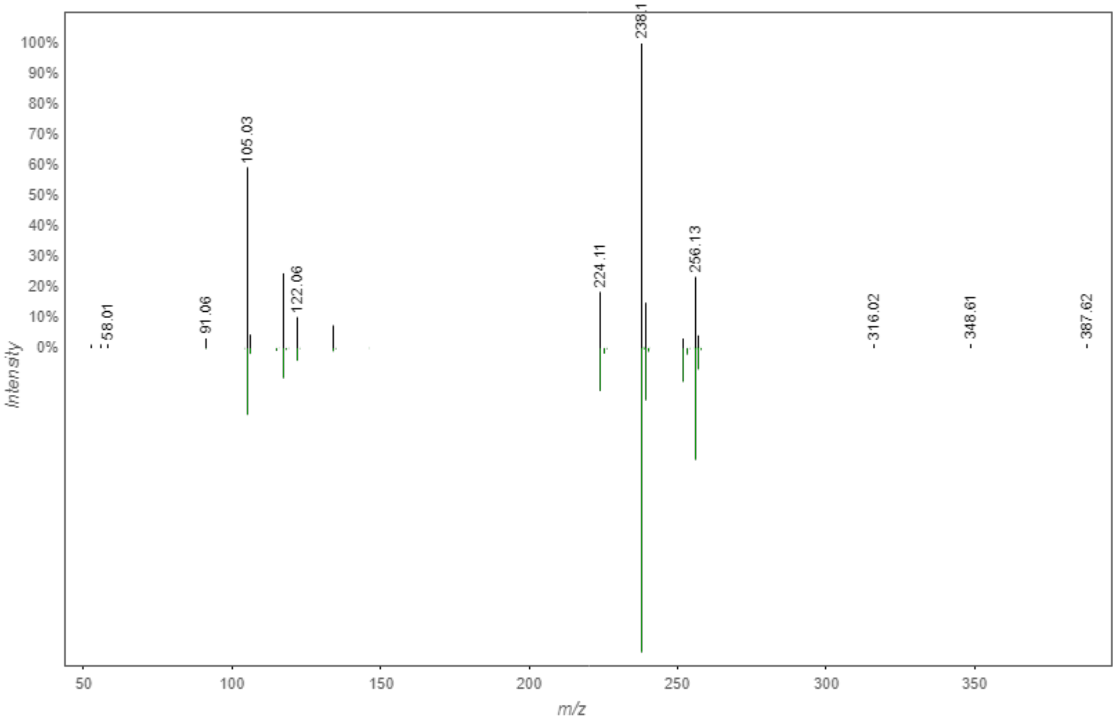


Figure S11. GNPS hit/MS data for asperphenamate.

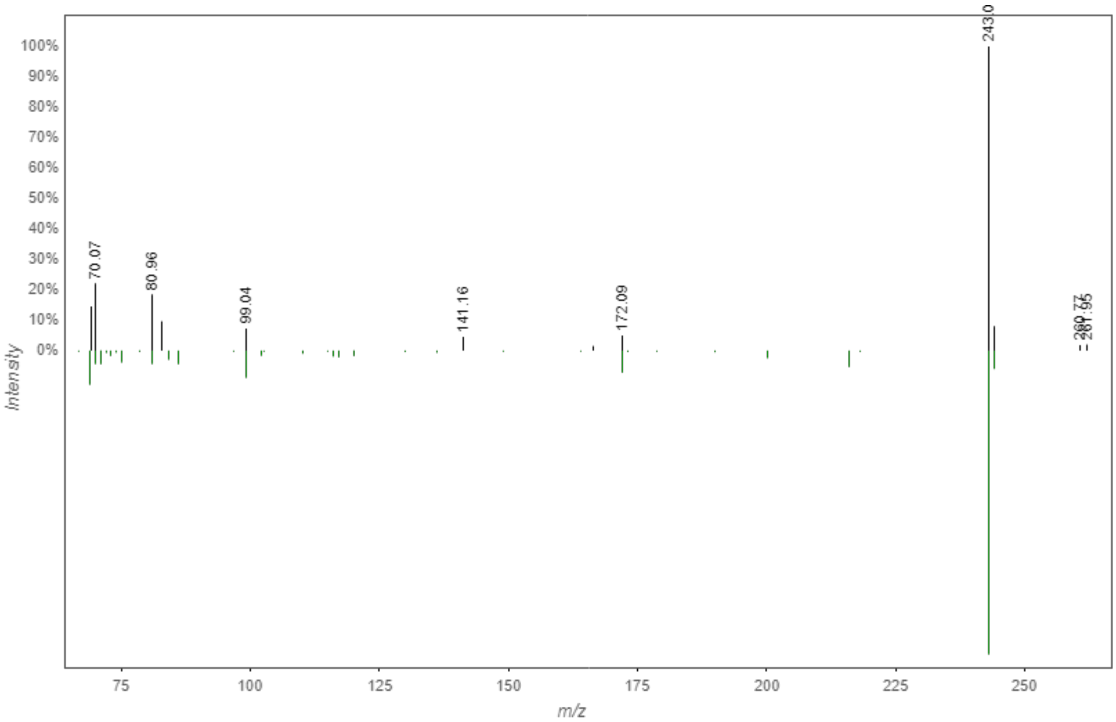

Figure S12. GNPS hit/MS data for riboflavin.

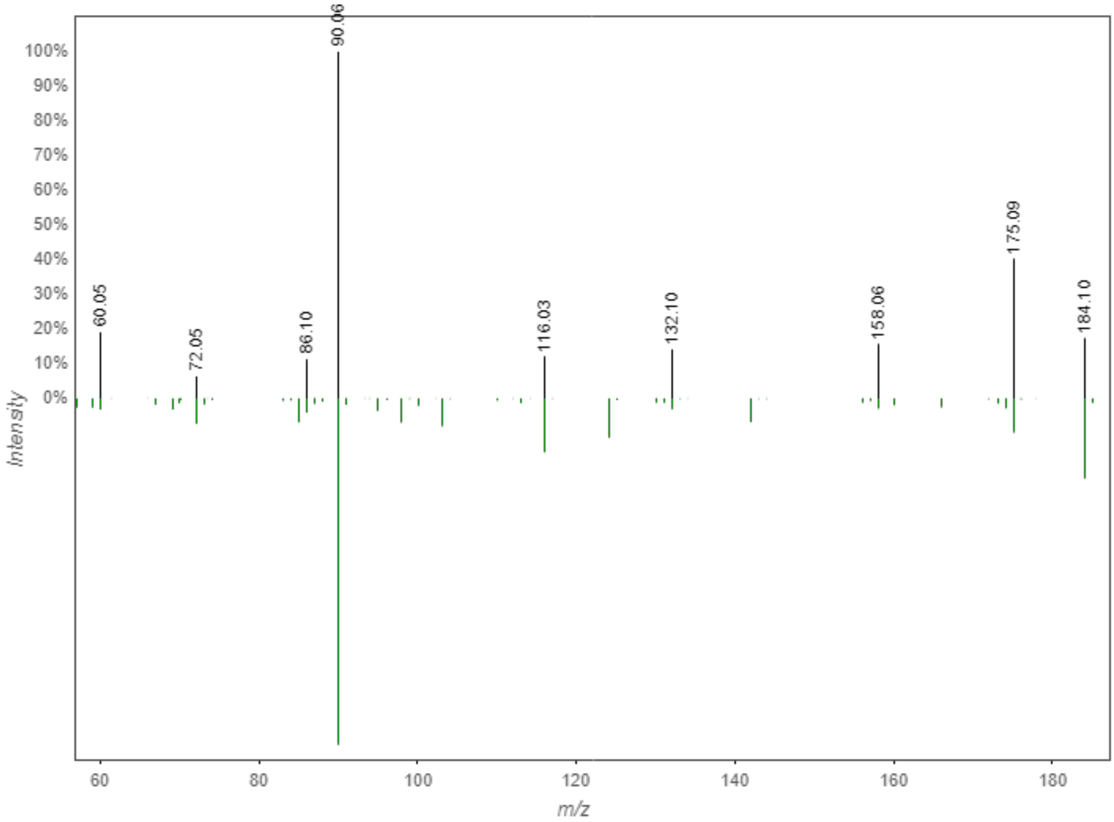

Figure S13. GNPS hit/MS data for pantothenic acid.
